# SERCA2 regulates proinsulin processing and processing enzyme maturation in the pancreatic β cell

**DOI:** 10.1101/2022.06.13.495980

**Authors:** Hitoshi Iida, Tatsuyoshi Kono, Chih-Chun Lee, Preethi Krishnan, Matthew C. Arvin, Staci A. Weaver, Timothy S. Jarvela, Robert N. Bone, Xin Tong, Peter Arvan, Iris Lindberg, Carmella Evans-Molina

**Author notes:** Address correspondence and requests for reprints to: Carmella Evans-Molina, MD, PhD, Indiana University School of Medicine, 635 Barnhill Drive, MS 2031A, Indianapolis, IN 46202. These authors contributed equally to this work (Hitoshi Iida, Tatsuyoshi Kono).

## Abstract

Increased circulating levels of incompletely processed insulin (i.e. proinsulin) are observed clinically in both type 1 and type 2 diabetes; however, the mechanisms underlying impaired proinsulin processing remain incompletely understood. Here, we identify the sarcoendoplasmic reticulum Ca^2+^ ATPase-2 (SERCA2) pump and β cell ER Ca^2+^ as key regulators of systemic glucose tolerance and proinsulin processing. We generated mice with a β cell-specific SERCA2 deletion (βS2KO) and SERCA2 deficient INS-1 cells to show that SERCA2 loss increases systemic and pancreatic levels of proinsulin protein and leads to aberrant localization of proinsulin within the proximal β cell secretory pathway. These defects in proinsulin processing were linked to reduced maturation of the proinsulin processing enzymes PC1/3 and PC2, suggesting a model whereby chronic ER Ca^2+^ depletion in the β cell, which is observed in many pathological conditions, impairs the spatial regulation of prohormone trafficking, processing, and maturation within the β cell secretory pathway.

## INTRODUCTION

Loss of coordinated insulin production and secretion from pancreatic β cells is a key pathological feature of both type 1 (T1D) and type 2 diabetes (T2D). Insulin production is an energetically consuming process that involves translation of preproinsulin mRNA on the rough endoplasmic reticulum (ER) and co-translational insertion of the nascent preproinsulin molecule into the ER lumen, where the N-terminal signal peptide is cleaved to form proinsulin (Liu et al., 2014; Vasiljevic et al., 2020; Weiss et al., 2000). Following disulfide bond formation and terminal protein folding within the ER and Golgi complex, proinsulin is packaged into immature secretory granules and routed to the regulated secretory pathway within the trans-Golgi network (TGN) (Arvan and Halban, 2004). The final steps of proinsulin processing and maturation involve cleavage of intact proinsulin into mature insulin and C-peptide by the proteolytic enzymes prohormone convertase 1/3 (PC1/3 encoded by the *PCSK1* gene), PC2 (*PCSK2*), and carboxypeptidase E (*CPE*), in a sequential process that begins in the TGN and is completed within the granule lumen. Secretion of mature insulin and C-peptide in response to glucose and other nutrient cues is a multistage process in which insulin-containing vesicles are transported to the plasma membrane for priming, docking, and fusion during exocytosis (Liu et al., 2021).

Under a variety of stress and disease conditions including overnutrition or genetic predisposition, the protein processing capacity within the β cell secretory compartment is overwhelmed, leading to the accumulation of inadequately processed proinsulin (Bollheimer et al., 1998; Hostens et al., 1999; Roomp et al., 2017; Sims et al., 2019b). Importantly, aberrant accumulation of proinsulin relative to insulin is observed in islets within pancreatic tissue from human organ donors with T1D, T2D, and pre-diabetes (Rodriguez-Calvo et al., 2021; Rodriguez-Calvo et al., 2017; Sempoux et al., 2001; Sims *et al*., 2019b). Clinically, this defect in protein processing and secretion manifests as an elevated ratio of secreted proinsulin:C-peptide or proinsulin:insulin that can be detected in the serum or plasma prior to the onset of both T1D and T2D, with persistence of this phenotype in established disease (Breuer et al., 2010; Egan et al., 2021; Leete et al., 2020; Pfutzner et al., 2015; Sims et al., 2016; Tong et al., 2020; Watkins et al., 2016). We and others have shown impaired proinsulin processing in ex vivo models of diabetes and metabolic stress (Arunagiri et al., 2019; Bollheimer *et al*., 1998; Hostens *et al*., 1999; Roomp *et al*., 2017; Scheuner et al., 2005; Sims et al., 2019a); however, the molecular pathways responsible for defective proinsulin processing in vivo during the development of diabetes remain poorly understood

ER calcium (Ca^2+^) depletion is a common pathway underlying multiple metabolic stressors in both T1D and T2D. Under normal conditions, steady state ER Ca^2+^ levels are maintained through the balance of Ca^2+^ transport into the ER lumen by the sarco-endoplasmic reticulum Ca^2+^ ATPase (SERCA) pump and Ca^2+^ release via the inositol trisphosphate receptors (IP3Rs) and ryanodine receptors (RyRs) (Gilon et al., 2014; Luciani et al., 2009; Tong et al., 2016; Yamamoto et al., 2019). We have shown previously that SERCA2 expression and activity are reduced in mouse models of diabetes, and that mice with SERCA2 haploinsufficiency manifest impaired glucose tolerance, reduced insulin secretion, an elevated immature to mature insulin granule ratio within the β cells, and increased proinsulin secretion when challenged with a high fat diet (Evans-Molina et al., 2009; Kono et al., 2012; Tong *et al*., 2016).

To investigate how loss of SERCA2 and chronic ER Ca^2+^ depletion impact proinsulin processing, we generated β cell-specific SERCA2-null C57BL/6J mice (βS2KO) and utilized a SERCA2-null rat insulinoma cell line (S2KO). We show that βS2KO mice have age-dependent glucose intolerance, increased serum and pancreatic proinsulin/insulin levels, decreased ER Ca^2+^ levels, and reduced glucose-stimulated Ca^2+^ synchronicity in isolated islets. Moreover, βS2KO islets and S2KO cells exhibited reduced expression of the active forms of the proinsulin processing enzymes, PC1/3 and PC2, and increased proinsulin accumulation within the proximal secretory pathway. Taken together, these findings identify a previously unappreciated role for SERCA2 and ER Ca^2+^ in the spatial regulation of prohormone trafficking, processing, and activation within the β secretory pathway.

## RESULTS

### Pancreatic β cell-specific SERCA2 deletion results in age-dependent glucose intolerance and impaired insulin secretion

Over 14 different isoforms of SERCA have been described to have tissue specific expression and alternative mRNA splicing (Periasamy and Kalyanasundaram, 2007). Within the β cell, SERCA2 and SERCA3 are both expressed, and we have identified SERCA2b as the predominant isoform (Kono *et al*., 2012). Therefore, to determine the role of SERCA2 in β cell biology, pancreatic β cell-specific SERCA2 knockout mice (βS2KO) were generated by crossing Ins1^cre/+^ and SERCA2^flox/flox^ mice (Thorens et al., 2015). The efficiency of SERCA2 deletion in isolated islets was approximately 95% at the protein level (Figures 1A-B) and 85% at the mRNA level (Figure 1C). Protein and mRNA expression of SERCA2 remained unchanged in other tissues including the hypothalamus, heart, liver, and skeletal muscle (Figure 1A-C). βS2KO mice exhibited normal body weight gain, random glucose levels, and lean mass on a chow diet, whereas fat mass was slightly increased compared to controls (Supplemental Figure S1A-D). While 8-week old βS2KO male mice showed normal glucose tolerance when challenged with an intraperitoneal (IP) glucose injection (Figure 1D), they developed significant glucose intolerance by 24 weeks of age (Figure 1E). There was no phenotype in female mice at the same age (Supplemental Figure S1E-H). The glucose intolerance phenotype of 24-week old male βS2KO mice was not accompanied by changes in systemic insulin sensitivity (Figure 1F), suggesting reduced β cell function in βS2KO mice. In support of this conclusion, the serum insulin response to IP glucose injection was significantly lower in βS2KO mice compared to littermate controls (Figure 1G), with a trend towards reduced β cell mass (classically defined by anti-insulin immuno-positivity) in βS2KO compared to control mice (0.75 mg vs. 0.98 mg, p=0.0797) at 24 weeks of age (Figure 1H).

**Figure 1.**
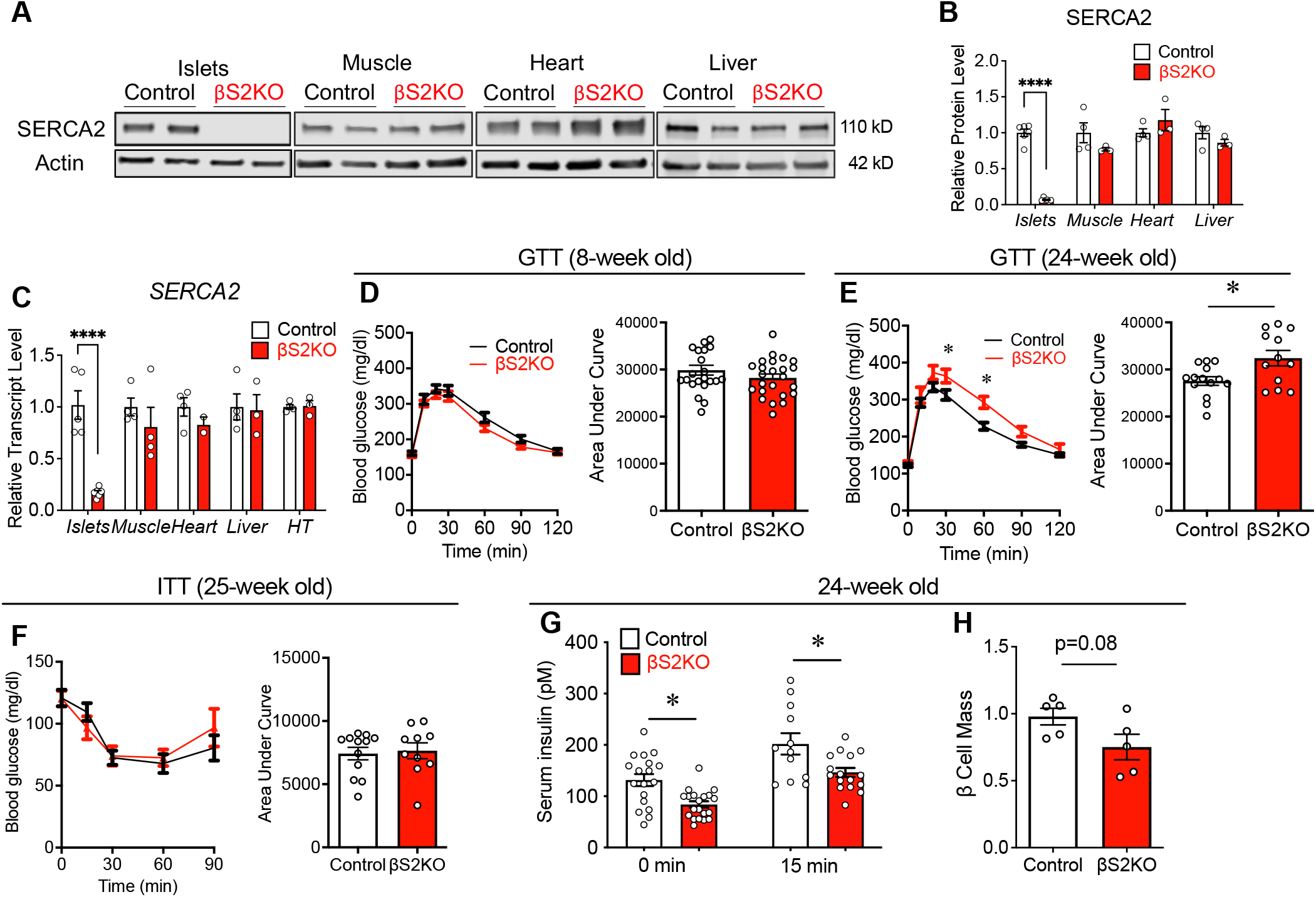
β cell-specific SERCA2 KO (βS2KO) mice exhibit age-dependent glucose intolerance and impaired insulin secretion without changes in insulin sensitivity. β cell-specific SERCA2 KO (βS2KO) and control SERCA2_flox/flox_ mice (Control) were fed a normal chow diet for 25 weeks. (A) Representative immunoblot performed using SERCA2 and actin antibodies in tissues from control and βS2KO mice. (B) Quantitation of immunoblotting results. Expression of SERCA2 was normalized to actin; n=3-6. (C) SERCA2 (*Atp2a2)* transcript levels in tissues of control and βS2KO mice were determined by RT-qPCR; n=3-6. HT: Hypothalamus. (D-E) Intraperitoneal glucose tolerance tests (GTT) were performed in male control and βS2KO mice. Glucose (2 g/kg body weight) was administered after 6 h of fasting. The tests were performed at 8 (D) and 24 (E) weeks of age. Bar graphs show quantification of the Area Under Curve; n=12-24. (F) Insulin tolerance tests (ITT) were performed in male control and βS2KO mice at 25 weeks of age following injection of Regular insulin at a dose of 0.5 mIU/g body weight. Bar graph shows AUC quantification; n=10-11. (G) In vivo glucose-stimulated insulin secretion (GSIS) assays were performed in male control and βS2KO mice after 6 h of fasting and administration of glucose (2 g/kg body weight). Insulin levels were measured by ELISA at baseline and 15 minutes after glucose injection. n=12-15. (H) Pancreatic β cell mass was assessed by insulin immunostaining in the pancreas of control and βS2KO mice, n=5. Results are presented as the mean ± S.E.M. Replicate samples are indicated by open circles. Comparisons between two groups were performed using unpaired Student’s *t-*test. Multiple comparisons were made using a two-way ANOVA and Sidak’s post test.*Indicates statistically significant difference from control (*p<0.05, ****p<0.0001).

### β cell-specific SERCA2 deletion reduces ER Ca^2+^ levels and alters nutrient-induced Ca^2+^ signaling

To test whether SERCA2 deletion was sufficient to reduce ER Ca^2+^ levels, islets isolated from control and βS2KO mice were transduced with an adenovirus expressing the D4ER Cameleon probe under transcriptional control of the rat insulin promoter (Ravier et al., 2011). ER Ca^2+^ levels were then measured by Fluorescence Resonance Energy Transfer (FRET). Islets isolated from βS2KO mice demonstrated a significant reduction in ER Ca^2+^ levels compared to control islets (Figure 2A). Next, glucose-stimulated Ca^2+^ responses were assessed using the cytosolic Ca^2+^ dye, Fura-2AM (Figure 2B). Islets from βS2KO mice showed a slight but statistically significant reduction in baseline cytosolic Ca^2+^ levels (3.3% reduction vs. control, Figure 2C), equivalent phase 1 amplitude (Figures 2D), increased phase 1 duration (26.3% increase vs control, Figure 2E), reduced amplitude of the 2^nd^ phase response (39.7% reduction vs. control, Figure F), and a longer oscillatory period (80% increase vs. control, Figure 2G). To measure glucose-induced Ca^2+^ oscillations in individual β cells, isolated islets were transduced with an adenovirus expressing the cytosolic calcium indicator GCaMP6 under control of the rat insulin promoter. Notably, β cells within islets of βS2KO mice showed diminished synchronicity of Ca^2+^ oscillations, as shown by a significantly higher GCaMP6 signal variance among the β cell population (Figure 2H-K).

**Figure 2.**
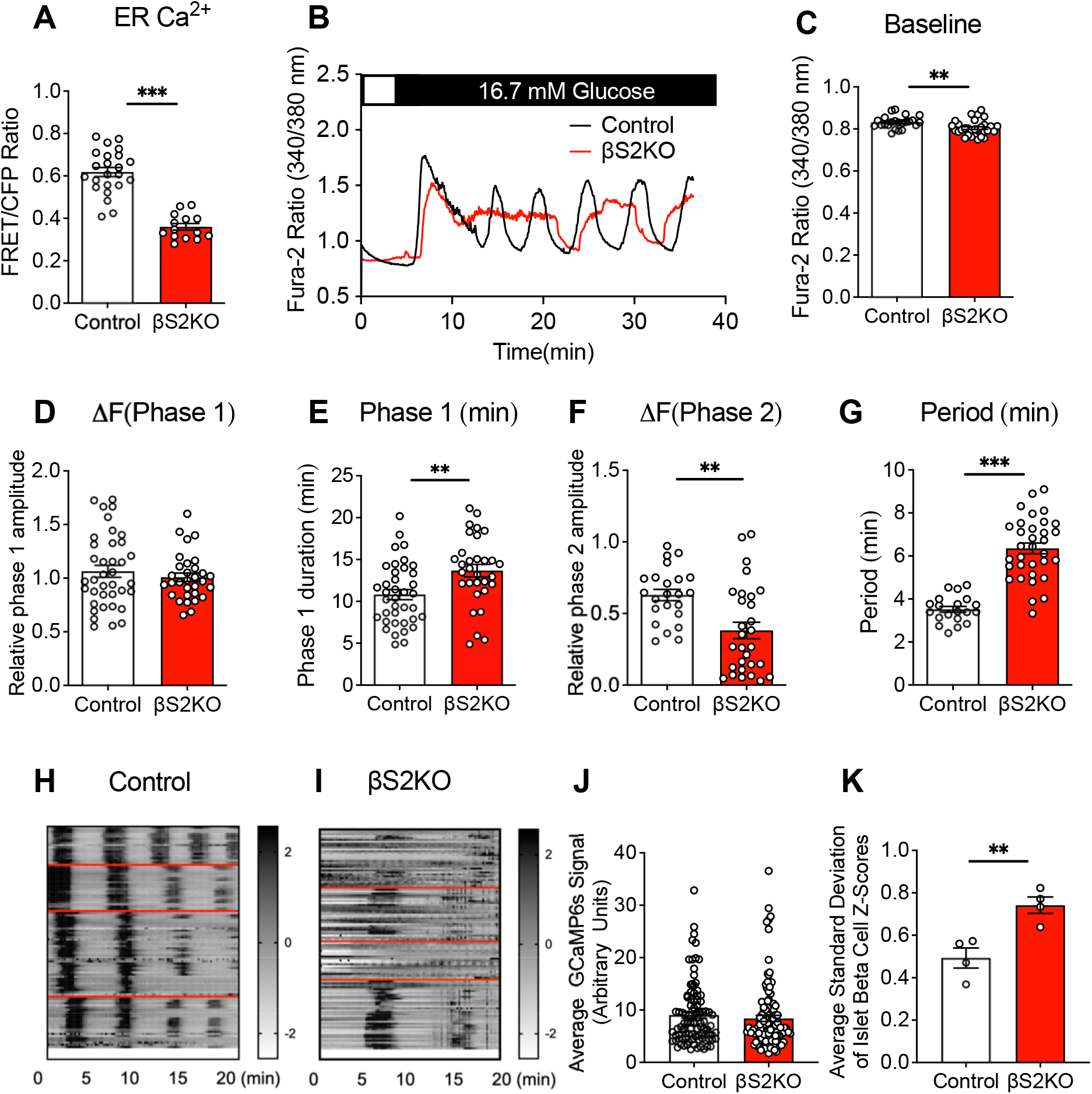
Islets from βS2KO mice exhibit reduced ER Ca_2+_ levels and altered glucose-stimulated Ca_2+_ oscillations. (A) Control and βS2KO islets were transduced with an adenovirus expressing the D4ER Cameleon probe under control of the rat insulin promoter. ER Ca_2+_ levels are expressed as the FRET/CFP ratio; n=14-23 islets from 3 mice. (B-G) Control and βS2KO islets were loaded with Fura-2 AM and Ca_2+_ imaging was performed. Representative recordings of cytosolic Ca_2+_ after stimulation with 16.7 mmol/L glucose (B). Quantification of the baseline (C), average phase 1 amplitude (D), phase 1 duration (E), phase 2 (oscillation) amplitude (F), and oscillation period (G); n=20-38 islets from 3 biological replicates. (H-K) Control and βS2KO islets were transduced with an adenovirus expressing GCaMP6s. (H and I) Example traces of phase 2 responses of GCaMP6 expressing islet cells treated with 16.7 mmol/L glucose. Regions of interest (ROI) were manually assigned to individual cells positive for GCaMP6s. The Z-score was calculated for each time point and each β cell as described in methods. Oscillations in cytosolic Ca_2+_ were illustrated by displaying the calculated Z-score of each β cell separated by islet with a red divider. Index bars on the right of the spectrograms showing the z-scores in grayscale. (J) The average GCaMP6s signal of control (n=98) and βS2KO (n=112) β cells was calculated and plotted. (K) Synchronicity was assessed by calculating the average standard deviation of islet β cells by taking the average of the standard deviation of Z-scores for each islet (n=4 per group) over the time course of the experiment. Results are presented as the mean ± S.E.M. Replicate samples are indicated by open circles. *p<0.05, **p<0.01, ***p<0.001 compared to control islets by a Student’s *t*-test.

### SERCA2 deficiency impairs proinsulin processing and processing enzyme maturation

To further investigate the mechanisms underlying the glucose intolerant phenotype observed in βS2KO mice, serum insulin and proinsulin levels were measured. Serum insulin levels were significantly decreased, coupled with an increase in serum proinsulin concentrations and an increase in the serum proinsulin:insulin (PI/I) ratio in βS2KO mice (Figures 3A-C). Consistent with these results, insulin content in the whole pancreas of βS2KO mice was decreased by ∼25%, while the proinsulin content and the pancreas PI/I ratio were both increased 1.5 to 2-fold (Figures 3D-F).

**Figure 3.**
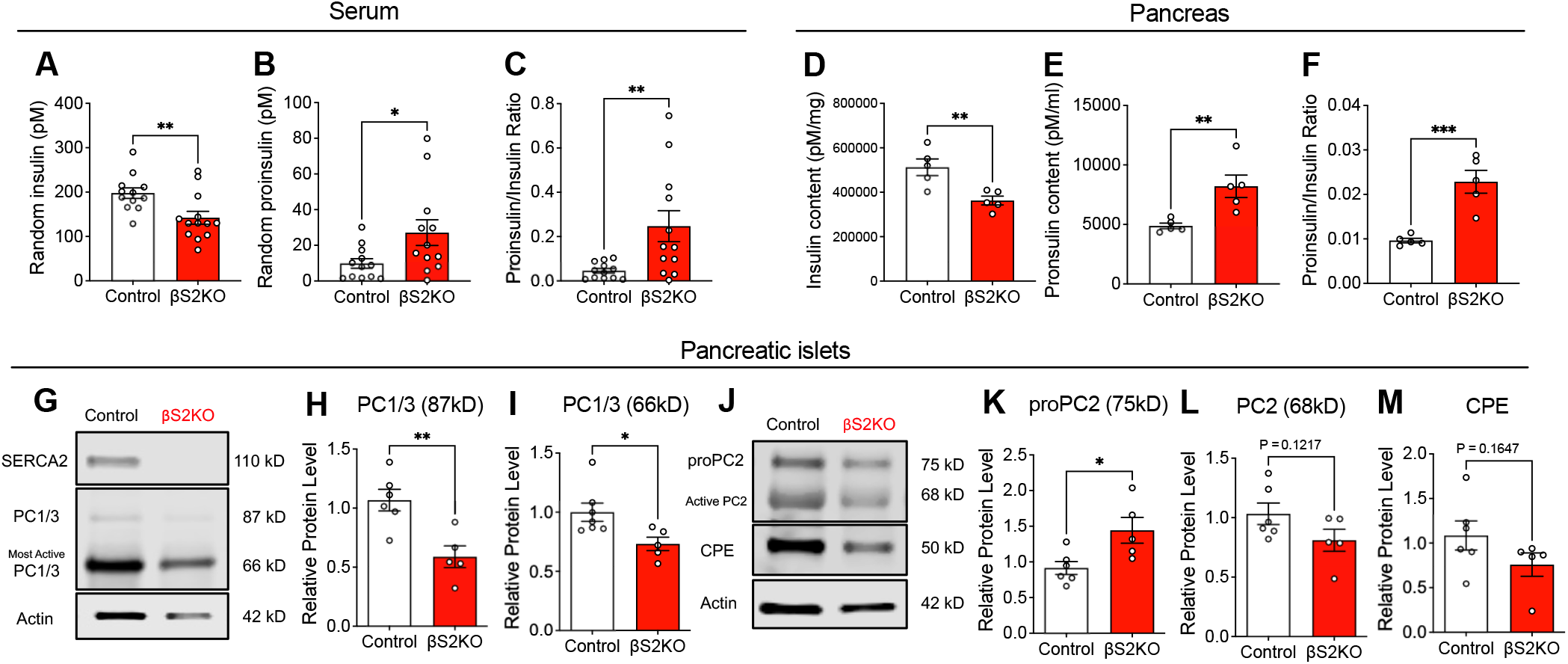
β cell-specific SERCA2 deficiency results in an increase of serum and total pancreas proinsulin/insulin ratios and alters the expression of prohormone convertases isoforms in islets. (A-C) Blood was collected from randomly fed control and βS2KO mice at 25 weeks of age and serum was isolated. Serum levels of insulin and proinsulin were determined by ELISA, and the proinsulin to insulin ratio was calculated; n=12-13 mice. (D-F) Total protein was extracted from the whole pancreas, levels of insulin and proinsulin were determined by ELISA, and the proinsulin to insulin ratio was determined; n=6 mice. (G and J) Representative immunoblots performed using antibodies against SERCA2, PC1/3, PC2, and CPE in islets obtained from 24-week old control and βS2KO mice. (H, I, K-M) Quantitation of PC1/3 (H and I), proPC2 (K), PC2 (L), and CPE (M) immunoblotting; results were normalized to actin expression; n=4-6. Results are presented as the mean ± S.E.M. Replicate samples are indicated by open circles. *Indicates statistically significant difference from control (*p<0.05, **p<0.01, ***p<0.001) by a Student’s *t*-test.

Given the prominent defect in proinsulin processing observed in our model, we assayed the effect of β cell-specific SERCA2 deletion on the expression of the three main proinsulin processing enzymes, PC1/3, PC2, and CPE. Normally, PC1/3, PC2, and CPE are synthesized as proenzymes and undergo sequential maturation and activation by autocleavage in different domains of the secretory pathway. Notably, production of both the intermediate/less active (87 kDa) and most active (66 kDa) forms of PC1/3 was significantly reduced, while proPC2 (75 kDa) was accumulated in βS2KO islets (Figures 3G-3K). There was a tendency towards reduced expression of the active form of PC2 (68 kDa) and CPE in βS2KO islets (Figures 3L-3M). Importantly, RT-qPCR analysis of islets showed no change in the mRNA levels of PC1/3, PC2, or CPE (Supplemental Figure S2A-E). Together, these results suggest that SERCA2 deletion in β cells impairs insulin biosynthesis at least in part by decreasing the levels of the most bioactive forms of the proinsulin endoproteolytic processing enzyme PC1/3 and PC2.

### Prohormone convertase enzyme activity is regulated in a SERCA2-dependent manner in β cell lines

To confirm a cell-autonomous relationship between SERCA2 and insulin processing, we analyzed prohormone convertase expression patterns in wild-type (WT) and SERCA2 deficient INS-1 cell lines [S2KO; (Tong *et al*., 2016)]. Similar to the results observed in islets from βS2KO mice (Figure 3), S2KO cells had reduced protein levels of the active forms of PC1/3 (66 kDa) and PC2 (68 kDa), with a tendency towards an increase in the PI/I ratio (Figure 4A-E). Adenoviral re-expression of SERCA2b in S2KO cells restored protein levels of the active forms of PC1/3, PC2, and CPE and decreased the PI/I ratio to levels seen in WT cells (Figure 4A-E). Fluorogenic enzyme assays showed an 81% reduction in PC1/3 enzyme activity and a 55% reduction in PC2 enzyme activity in S2KO cells compared to WT (Figure 4F-G), while the activity of both enzymes was partially rescued by adenoviral restoration of SERCA2 expression in S2KO cells (Figure 4F-G). Our results suggest a SERCA2-dependent regulation of proenzyme maturation and activation in the secretory pathway of pancreatic β cells (Figure 4H).

**Figure 4.**
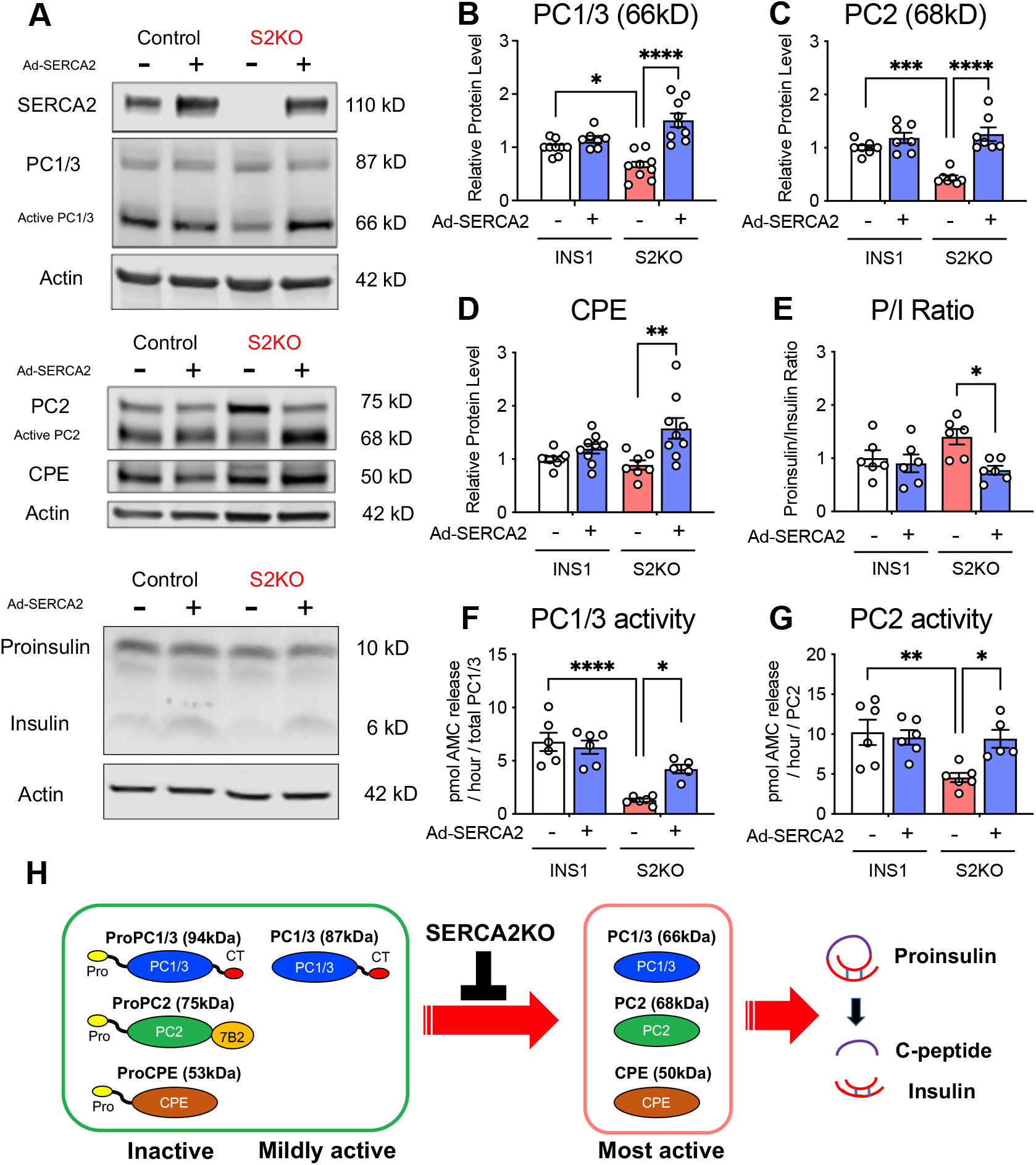
Enzyme activity of prohormone convertases is regulated in a SERCA2-dependent manner in β cell lines. (A) Representative immunoblots of SERCA2, PC1/3, PC2, CPE, proinsulin, and insulin in WT and S2KO cells transduced with empty adenovirus or an adenovirus expressing human SERCA2b-adenovirus (Ad-SERCA2). (B-E) Quantitation of immunoblotting results. Expression of proteins was normalized to actin expression; n=6-9. (F and G) WT and S2KO INS-1 cells were transduced with empty adenovirus or an adenovirus expressing human SERCA2b, and the enzyme activities of secreted PC1/3 and PC2 were determined using fluorometric assays. Enzyme activity was normalized to the protein amount of each enzyme, as determined by immunoblotting. n = 5-6 Results are presented as the mean ± S.E.M. Replicate samples are indicated by open circles. (H) Schematic illustration of the SERCA2KO function in maturation of prohormone convertases and insulin processing. Before becoming enzymatically capable of cleaving their targets including proinsulin, precursors to PC1/3, PC2, and CPE enzymes must be activated through a series of autoproteolytic events, most of which are pH- and calcium-dependent. *Indicates statistically significant difference (*p<0.05, **p<0.01, ***p<0.001, ****p<0.0001) by a two-way ANOVA and Sidak’s post test.

### RNA sequencing shows functional enrichment of genes involved in calcium signaling, diabetes pathophysiology, and secretory pathway function in islets from βS2KO mice

To gain insight into novel pathways through which β cell SERCA2 modified systemic glucose tolerance and proinsulin processing, RNA sequencing was performed using islets isolated from 17-week-old control and βS2KO male mice—a time-period prior to the development of glucose intolerance. We identified 453 differentially expressed mRNAs, of which 22 were upregulated and 431 (95%) were downregulated in βS2KO mice (Figure 5A; Supplementary Table S3). There was a tendency towards an increase in *Ins1* and a decrease in *Ins2* in the RNA seq data, and the real-time qRT-PCR showed no significant changes of *Ins1* or *Ins2* in βS2KO islets at 24-weeks old (Supplemental Figure S3). Pathway analysis identified 162 pathways that were significantly modulated and, after removing terms less relevant to our study, 112 pathways were retained (Supplementary Table S4). Gene Ontology (GO) terms were identified using Metascape, and terms with p<0.05 were considered significant (Figure 5B; Supplementary Table S5). Significantly altered GO terms included processes involved in calcium signaling and pathways relevant to diabetes pathophysiology (“apoptosis”, “antigen presentation”, and “oxidative stress”), with a notable enrichment of terms related to secretory function (“exocytosis”, “protein secretion”, “processing”, and “vesicle mediated transport”).

**Figure 5.**
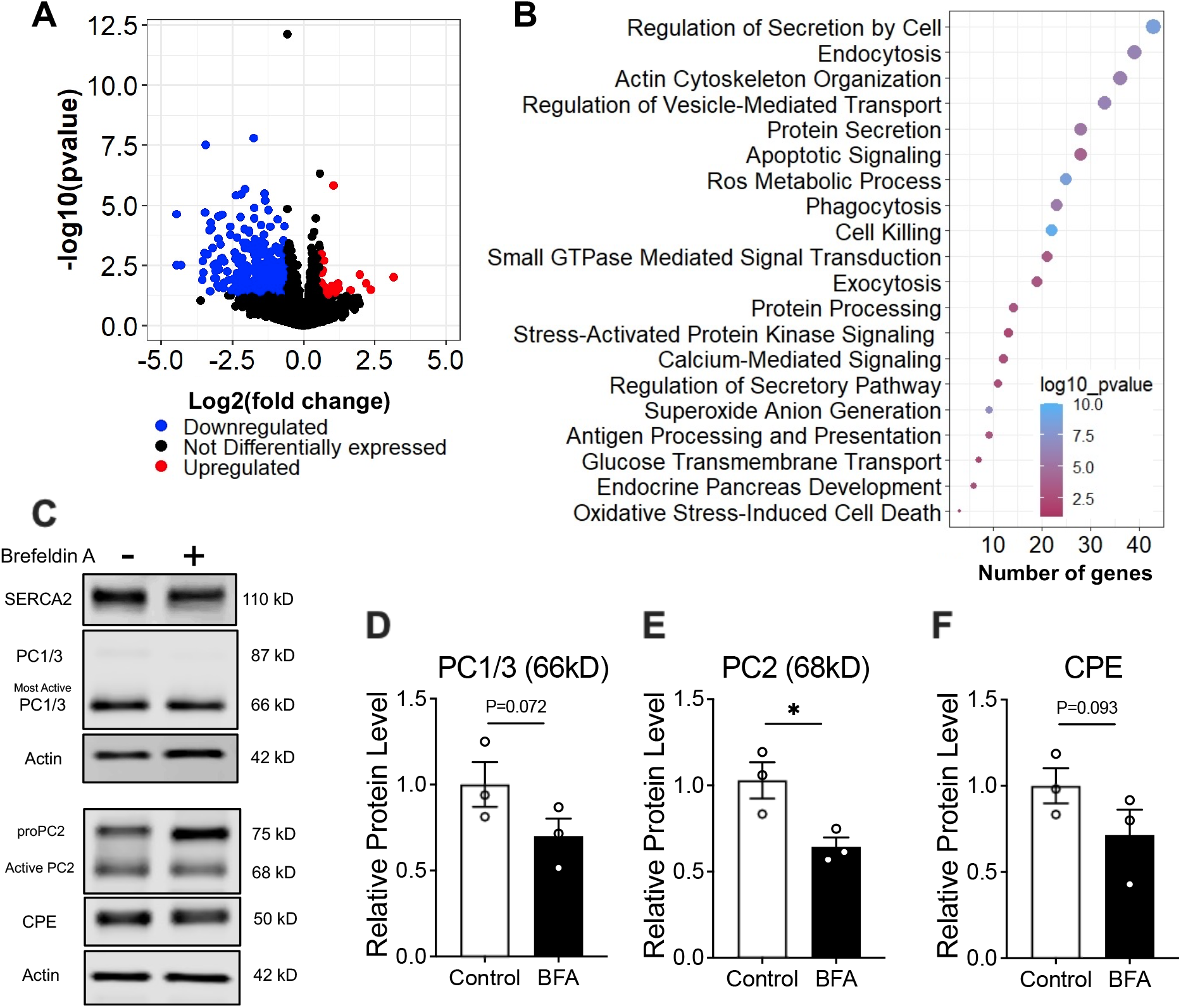
Bulk RNA sequencing and chemical inhibition of vesicle trafficking demonstrate a role for SERCA2 in prohormone maturation. (A-B) RNA isolated from control and βS2KO islets was subjected to bulk RNA sequencing analysis at 17-weeks of age. (A) Volcano plot indicating differentially expressed genes (FC ≥ 1.5, p < 0.05). Red indicates upregulated genes, blue indicates downregulated genes, and black indicates no change in expression in βS2KO islets compared to control islets. (B) Dot plot representing 20 Gene Ontology terms identified using the ggplot2 package in R. Size of the dot indicates the number of genes in each term. Enrichment analysis was performed using Metascape. (C-F) Isolated WT mouse islets were treated with 4 µM brefeldin A for 6 hours. (C) Representative immunoblots of PC1/3, PC2, and CPE. (D-F) Quantitation of immunoblotting results. Expression of proteins was normalized to actin; n=3. *Indicates statistically significant difference from control (p<0.05) by Student’s *t*-test.

### SERCA2 loss disrupts protein trafficking within the β cell secretory pathway

Enrichment for terms related to secretory pathway transport within the RNA sequencing dataset was notable given our observations of reduced proinsulin processing and impaired maturation of proinsulin processing enzymes, both of which are spatially regulated within different domains of the β cell secretory pathway. Thus, we hypothesized that chronic ER Ca^2+^ depletion within the β cell secretory pathway may cause a proinsulin processing defect via impaired anterograde protein trafficking. With this in mind, we treated WT mouse islets with brefeldin A (BFA), an inhibitor of anterograde protein transport. Similar to what we observed in SERCA2-deficient islets and cells, BFA treatment decreased active PC2 levels (68 kDa) and led to a trend towards a reduction in the most active forms of PC1/3 (66 kDa, 30.0% reduction in BFA-treated islets, p=0.072) and as well as a decrease in mature 50 kDa CPE (28.5% reduction in BFA-treated islets, p=0.093) (Figures 5C-F).

Consistent with the notion that chronic ER stress impairs anterograde trafficking and processing of protein cargo, we observed increased stability of the intermediate form of PC1/3 (87 kDa) and proPC2 (75 kDa) in S2KO cells (Figures 6A-C). The stability of these two proteins was prolonged further by treatment with the proteasome inhibitor MG132 (Figure 6A-C). Lastly, immunofluorescent staining was performed to investigate changes in protein localization in islets from βS2KO and control mice. Proinsulin staining intensity was significantly increased in islets of βS2KO mice (Figures 6D-E). Notably, in islets from βS2KO mice there was increased colocalization of proinsulin and lectin mannose-binding 1 (LMAN1), a marker of the intermediate region between the ER and the Golgi (ERGIC), as well increased colocalization of proinsulin and giantin, a marker of the Golgi apparatus (Figure 6F-G). These findings indicate that SERCA2 deficiency leads to an accumulation of proinsulin in the ERGIC and/or cis-Golgi complex in β cells, suggesting a defect in trafficking of protein cargo to more distal portions of the secretory pathway where normal processing occurs to produce mature insulin.

**Fig 6.**
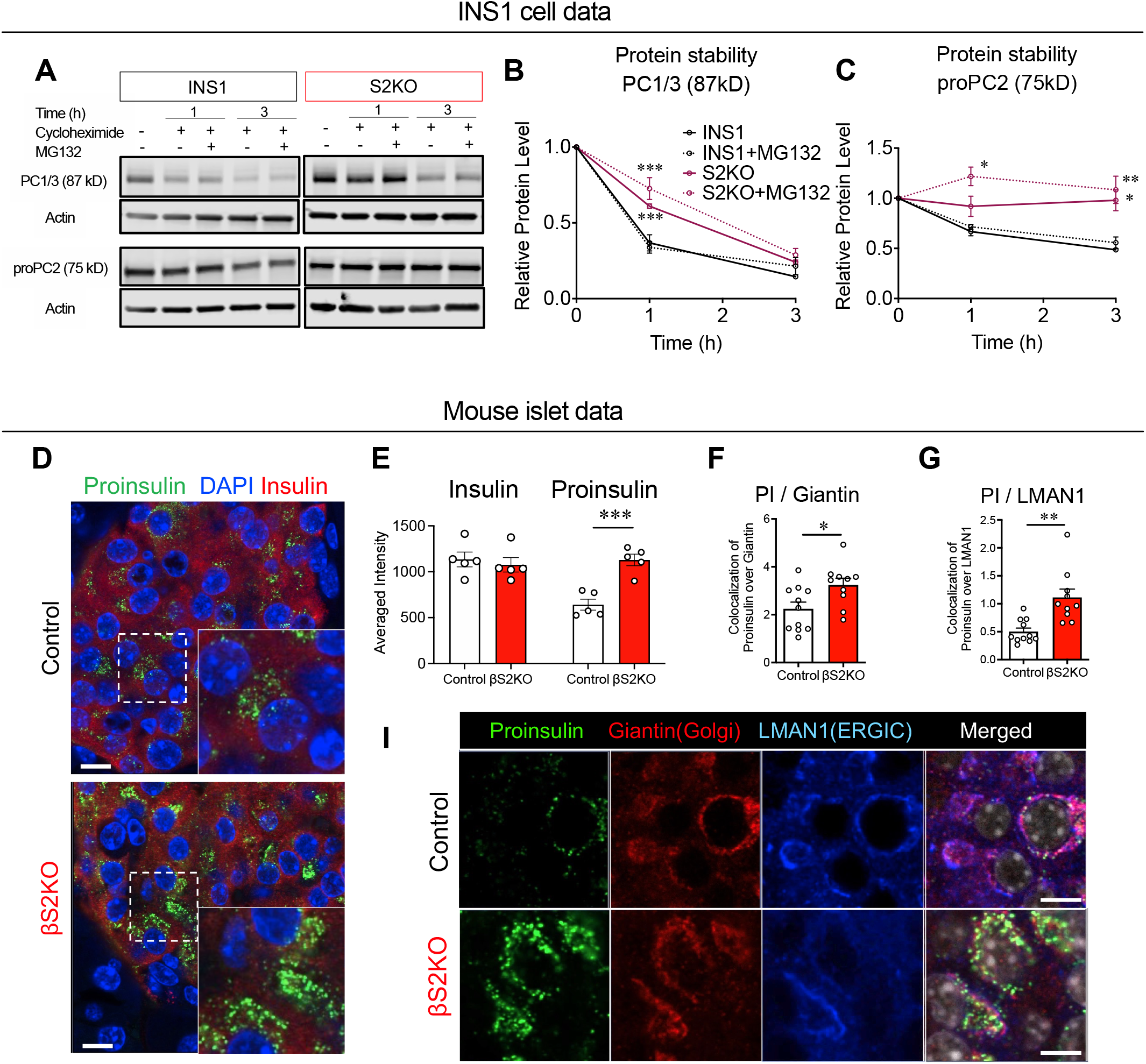
SERCA2 loss disrupts protein trafficking within the β cell secretory pathway. (A-C) WT and S2KO INS-1 cells were treated with 10 µM cycloheximide with or without 10 µM MG-132 for 9 hours prior to protein extraction. Representative immunoblots of the 87 kDa less active form of PC1/3 and proPC2 (A). Quantification of time-dependent changes in the level of 87 kDa PC1/3 (B) and 75 kDa proPC2 (C) after the addition of cycloheximide; n=3 independent experiments. (D-E) Pancreatic sections from control and βS2KO mice were stained for proinsulin, insulin, and DAPI. Representative immunofluorescent images (D) and quantification of protein expression levels using Image J (E). (F-G) Pancreatic sections from control and βS2KO mice were stained for proinsulin, giantin, and LMAN1. Quantification of protein expression levels and localization using Cell Profiler (F-G). Results are presented as the mean ± S.E.M. Replicate samples are indicated by open circles. *p<0.05, **p<0.01, ***p<0.001 vs. indicated groups. Two-way ANOVA was used for B and C, and Student’s *t*-test for E, F, and G.

### Glucolipotoxicity decreases active prohormone convertase isoform protein expression in mouse and human islets

A glucolipotoxic environment, similar to that observed in T2D, is known to inhibit insulin biogenesis (Bollheimer et al., 1998; Fontes et al., 2010), vesicle budding, protein trafficking, and ER lipid raft formation (Lytrivi et al., 2020; Preston et al., 2009). We have previously shown that glucolipotoxic treatment (GLT; 25 mM glucose and 0.5 mM palmitate) reduces SERCA2 expression and activity in mouse islets and INS-1 cells (Tong *et al*., 2016). To test the relevance of our findings in an in vitro model of T2D, we utilized GLT to evaluate changes in prohormone convertase expression in WT mouse islets and in human islets.

In mouse islets treated with GLT, SERCA2 protein expression was reduced by ∼80% and the intracellular PI/I ratio was increased (Figures 7A-E). In addition, GLT reduced the levels of the most active (66 kDa) form of PC1/3 but increased its intermediate form (87 kDa) (Figures 7F-G). Active PC2 levels showed a similar tendency in response to GLT treatment, but the changes did not reach statistical significance (Figures 7H-I), while the active form of CPE was significantly reduced by GLT treatment in mouse islets (Figure 7J). Finally, a similar immunoblot analysis was performed in islets from non-diabetic human donors treated with GLT. Overall, the active forms of PC1/3, PC2, and CPE were all significantly reduced in GLT-treated human islets (Figures 7K-P).

**Figure 7.**
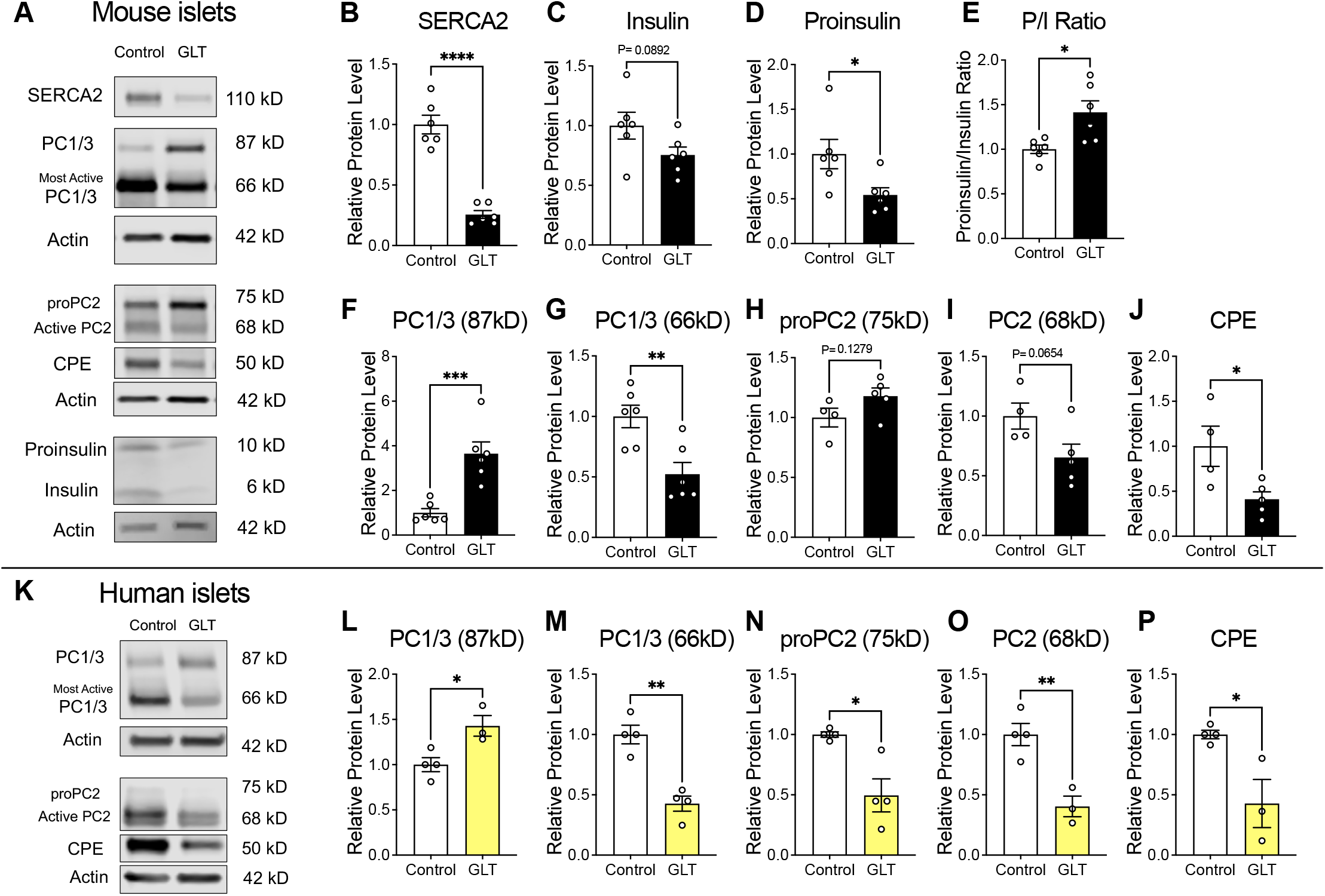
Decreased active prohormone convertase isoform expression in GLT-treated mouse islets and human islets. (A-J) Isolated WT mouse islets were subjected to GLT treatment (25mM glucose + 0.5 mM palmitate) for 24 hours. Representative immunoblots of SERCA2, proinsulin, insulin, PC1/3, PC2, and CPE (A) and quantitation of immunoblotting results (B-J). Expression of proteins was normalized to actin; n=4-6. (K-P) Human islets from non-diabetic donors were subjected to GLT treatment (25mM glucose + 0.5 mM palmitate) for 24 hours. Representative immunoblots of PC1/3, PC2, and CPE (K) and quantitation of immunoblotting results (L-P). Expression of proteins was normalized to actin; n=3-4. Results are presented as the mean ± S.E.M. Replicate samples are indicated by open circles. *Indicates statistically significant difference from control (*p<0.05, **p<0.01, ***p<0.001, ****p<0.0001) by a Student’s *t*-test.

## DISCUSSION

Here, we developed mice with a β cell-specific SERCA2 deletion (βS2KO) to determine the impact on ER Ca^2+^ and global pancreatic β cell function. Our results show that loss of SERCA2 was sufficient to reduce steady state β cell ER Ca^2+^ levels, to impair glucose-stimulated Ca^2+^ responses, and to reduce synchronization between β cells within islets. In addition, βS2KO mice exhibited mild age-dependent glucose intolerance and reduced in vivo glucose-stimulated insulin secretion. This overall phenotype is consistent with our previous study of high fat diet fed mice with global SERCA2 haploinsufficiency (Tong *et al*., 2016). However, the β cell-specific KO reported here allows us to attribute defects in metabolic function to SERCA2 activity specifically in the β cell. Overall, our results underscore the importance of the ER Ca^2+^ pool in maintaining normal β cell function. Moreover, our findings are consistent with other studies that have documented the importance of SERCA2 and ER Ca^2+^ homeostasis in other disease states, including dermatologic disorders (Celli et al., 2012) and Alzheimer’s disease (Britzolaki et al., 2018).

Notably, we found that SERCA2 deletion in the β cell led to a prominent defect in proinsulin processing that was reflected in both the pancreas and serum of βS2KO mice. Defects in proinsulin processing have been observed in both T1D and T2D (Sims *et al*., 2019a), and activation of a variety of β cell stress pathways, including endoplasmic reticulum, oxidative, and lipotoxic stress, have been associated with ER Ca^2+^ depletion and dysfunctional proinsulin processing (Biden et al., 2014; Cnop et al., 2005; Han et al., 2015). At present, the specific molecular mechanisms underlying defects in proinsulin processing have not been well characterized. Previous studies have demonstrated reduced mRNA expression of PC1/3, PC2, and CPE in human islets treated with proinflammatory cytokines and other metabolic stressors (Sims *et al*., 2019b). However, in the work reported here, mRNA expression levels of the major processing enzymes were unaffected in βS2KO islets. Instead, our data indicate distinct changes in the post-transcriptional processing of these enzymes. Under normal conditions, PC1/3, PC2, and CPE are synthesized as proenzymes and undergo sequential maturation by autocleavage within different domains of the secretory pathway. This process is closely regulated by both pH and Ca^2+^ (illustrated in Figure 4H), thus providing spatial regulation of proenzyme activation and proinsulin processing (reviewed in (Seidah, 2011). In islets from βS2KO mice and S2KO INS-1 cells, we observed reductions in the most active forms of PC1/3, PC2, and CPE and accumulation of the immature proenzyme proPC2. We showed that these changes were associated with reduced PC1/3 and PC2 enzyme activity, while SERCA2 adenoviral overexpression rescued enzyme activity and maturation.

Changes in enzyme maturation in islets from βS2KO mice and in S2KO INS-1 cells suggest a model whereby chronic ER Ca^2+^ depletion and SERCA2 deficiency disrupt forward trafficking of protein cargo in the β cell, leading to an altered distribution of these cargoes within the secretory pathway, which further contributes to impaired proenzyme maturation and prohormone processing. Several lines of evidence support this model. First, results from RNA sequencing performed using islets from male βS2KO mice showed downregulation of genes that were enriched for terms related to vesicle-mediated trafficking and secretory function. Second, brefeldin A, which inhibits ER to Golgi vesicle trafficking, partially recapitulated the impairments in PC1/3 and PC2 expression observed in our SERCA2 deficient models. Third, in S2KO INS-1 cells, we observed an increased half-life of both proPC2 and the less-active/intermediate form of PC1/3. Lastly, in pancreas sections from βS2KO mice, we demonstrate increased co-localization of proinsulin with markers of the early compartments of the secretory pathway, including the intermediate compartment between the ER and Golgi.

The mechanisms by which SERCA2 and ER Ca^2+^ regulate processing of hormone maturation and trafficking in the β cell remain a matter of some speculation. Ca^2+^ is a required cofactor for PC1/3 and PC2 activity [reviewed in (Chen et al., 2018)], so decreased secretory pathway Ca^2+^ could directly affect convertase activity on prohormone substrates, as well as indirectly impact prohormone maturation via reduced intermolecular cleavage of 87 kDa PC1/3 to the more active 66 kDa form (Zhou and Lindberg, 1994). However, independent of reduced availability of Ca^2+^ as an enzyme cofactor, we suggest that loss of SERCA2 also results in impaired or inefficient anterograde trafficking of both processing enzymes and proinsulin through the various compartments of the secretory pathway where these cargoes undergo their sequential maturation, processing, and activation.

The importance of ER-to-Golgi trafficking on the development of diabetes has been highlighted recently in a study of neonatal diabetes caused by mutation of *YIPF5* (De Franco et al., 2020), a protein that cycles between the ER and the Golgi and is predominantly localized in the ER and the ER-Golgi intermediate compartments. The deletion of *YIPF5* caused a 5.5-fold increase in proinsulin staining, a 70% reduction in insulin staining, and was associated with a newly described monogenic form of diabetes (De Franco *et al*., 2020). At least one prior study has linked lipotoxic stress with impaired ER-to-Golgi trafficking in the β cell (Boslem et al., 2011). This is notable because we have shown that glucolipotoxicity reduces SERCA2 expression (Tong *et al*., 2016), and treatment of human islets with glucolipotoxicity in the present study partially phenocopied the altered PC1/3 and PC2 expression patterns observed in our SERCA2-deficient mouse model. A recent report from Ramzy *et al*. demonstrated an unexplained accumulation of PC2 in islets from human organ donors with T2D (Ramzy et al., 2020). Our data extend these findings by showing a prolonged protein half-life for 87 kDa PC1/3 and proPC2 in SERCA2-deficient cells, which we hypothesize may arise from inefficient trafficking within the secretory pathway.

In this regard, luminal Ca^2+^ has been shown to affect ER to Golgi trafficking and COPII vesicle budding, delivery, and fusion with the Golgi in other cell types (Brandizzi and Barlowe, 2013). In keratinocytes, SERCA2 deficiency reduced ER to Golgi transport of desmosomal cadherin cargo (Li et al., 2017). Consistent with this observation, in individuals with Darier White disease, a rare genetic icthyosis disorder resulting from SERCA2 haploinsufficiency, lesion keratinocytes exhibit ER retention of cadherin, a transmembrane protein located at the plasma membrane that typically requires trafficking to the membrane (Celli *et al*., 2012). Only one group has examined proinsulin to insulin ratios in individuals with Darier White; they described a non-significant trend towards increased ratios in a small cohort of affected individuals compared to matched controls (Ahanian et al., 2020). This same group found that individuals with Darier White have an increased relative risk of developing T1D (Cederlöf et al., 2019).

There are some limitations of our study that should be acknowledged. Our findings could have relevance to other proteins and hormones that require processing in β cells and other secretory cell types, such as salivary glands, alpha cells, or intestinal cells that rely heavily on proper protein processing and vesicle trafficking. However, additional experiments are required to understand whether defects in trafficking are generalizable or cargo-specific, and whether impaired forward trafficking engages alternative pathways of cellular egress in the diabetic β cell. Additionally, during the course of our study, we uncovered a polymorphism in the *P2rx7* gene that impacts many 129-originating transgenic mouse lines (Er-Lukowiak et al., 2020). To account for this passenger mutation, all comparisons in this manuscript were made using Atp2a2^fl/fl^; Ins1^Cre^-negative mice as controls.

Notwithstanding these limitations, our results provide additional insight into the function of SERCA2 and the importance of luminal Ca^2+^ in the regulation of secretory pathway function in the pancreatic β cell, and illustrate a unique relationship between SERCA2 activity and the spatial regulation of proenzyme and prohormone processing in the β cell.

## MATERIALS AND METHODS

### RESOURCE AVAILABILITY

#### Lead Contact

Further information and requests for resources and/or reagents should be directed to and will be fulfilled by Dr. Carmella Evans-Molina.

#### Materials availability

This study did not generate new unique reagents.

#### Data and code availability

- The datasets supporting the current study are available from the lead contact on request. Sequencing data generated in this study will be deposited to the NCBI Gene Expression Omnibus repository upon paper acceptance.
- This paper does not report original code.
- Any additional information required to reanalyze the data reported in this paper is available from the lead contact upon request.

### EXPERIMENTAL MODEL AND SUBJECT DETAILS

#### Animals

Male B6(Cg)-Ins1^tm1.1(cre)Thor^/J mice (Ins1^cre^) on C57BL/6J background (MGI:5614359) were purchased from Jackson Laboratory. Atp2a2^tm1.1Iemr^ mice on 129P2/OlaHsd background (MGI:4415164) were a kind gift from Dr. Ole M. Sejersted (University of Oslo). Atp2a2^tm1.1Iemr^ mice were backcrossed for more than 10 generations onto a C57BL6/J background, and β cell-specific SERCA2 knockout (βS2KO) mice were generated by crossing Atp2a2^tm1.1Iemr^ mice on C57BL/6J background with Ins1^tm1.1(cre)Thor^ mice (MGI:5614359). Mice were studied between 4 and 25 weeks of age.

During our analysis of these mice, we became aware of a polymorphism in the *P2rx7* gene that impacts many 129-originating transgenic mouse lines. To account for this passenger mutation, all comparisons in this manuscript were made using Atp2a2^fl/fl^; Ins1 Cre negative mice as controls. Therefore, we are able to ascribe our observed phenotype directly to SERCA2 deletion, rather than to an off-target effect of the *P2rx7* passenger mutation inherent to the Atp2a2^tm1.1Iemr^ mice. The data on this polymorphism are available online (Er-Lukowiak *et al*., 2020).

Because female βS2KO mice showed normal glucose tolerance, islets from male mice were used for ex vivo analyses. Littermates were randomly assigned to experimental groups. Mice were group housed at 22 ± 2°C under a 12h:12h light-dark cycle (7 AM-7 PM), with *ad libitum* access to chow rodent diet (Teklad 2018SX). Health was monitored daily by veterinary staff at the IU School of Medicine, and all mice used for experiments showed normal health.

All protocols involving mice were approved by the Indiana University Institutional Animal Care and Use Committee.

#### Cell lines

The male INS-1 832/13 cell line is an established model for studying glucose-stimulated insulin biosynthesis and secretion of insulin (Hohmeier et al., 2000). The INS-1 cell line was generated by irradiation of a male rat insulinoma (Chick et al., 1977), and the INS-1 832/13 line was generated by transfecting INS-1 cells with a plasmid containing the human insulin gene under the cytomegalovirus promoter (Asfari et al., 1992). A SERCA2 KO INS-1 cell line (S2KO) was generated by the Genome Engineering Center (GEC) at Washington University (St. Louis, MO) from WT INS-1 832/13 cells using clustered regularly interspaced short palindromic repeats (CRISPR)/Cas 9 technology (Tong *et al*., 2016). S2KO and WT INS-1 cells were maintained at 37°C in an atmosphere supplemented with 5% CO_2_. The cells were grown in RPMI1640 (Gibco, Waltham, MA) containing 10% fetal bovine serum (FBS), 100 U/mL penicillin, 100 μg/mL streptomycin, 10 mM HEPES, 2 mM L-glutamine, 1 mM sodium pyruvate, and 50 μM β-mercaptoethanol.

#### Human Tissue

Human cadaveric islets were obtained from the Integrated Islet Distribution Program (IIDP, City of Hope Duarte, CA). Donor characteristics are shown in Supplemental Table S2. Immediately upon receipt, human pancreatic islets were placed in DMEM containing 5.5 mM glucose, 10% FBS, 100 U/mL penicillin, and 100 μg/mL streptomycin and incubated overnight at 37°C in an atmosphere containing 5% CO_2_ (Johnson et al., 2014). Human islets distributed by IIDP are from cadaveric organ donors from which at least one other organ has been approved for transplantation; tissues are considered “exempt” from Human Studies Approval.

### METHOD DETAILS

#### Mouse metabolic studies

Body weight and random blood glucose levels were measured every other week. Blood glucose levels were measured in whole blood collected from the tail vein using a Contour® glucometer. Glucose tolerance tests (GTT) were performed following intraperitoneal (IP) administration of 2 g/kg body weight glucose after 6 h of fasting, as previously published (Tong *et al*., 2016). D-(+)-glucose (Sigma) was prepared by adding 2.5 g powdered glucose to 10 mL sterile water, and letting it dissolve at 4°C overnight. Glucose-stimulated insulin secretion (GSIS) was determined *in vivo* by injection of glucose (2g/kg body weight, IP) after 6 h of fasting and collection of serum at 0 and 15 min after glucose injection. Insulin tolerance tests (ITT) were performed after 3 h of fasting followed by IP administration of 0.5 mIU/kg Regular insulin (Humulin R NDC-0002-8215-01). Insulin and proinsulin levels in serum and tissues were measured using ELISA kits. Manufacturer information noted that there is no cross reactivity with insulin or c-peptide for Rat/Mouse Proinsulin Elisa kit (Cat# 80-PINMS-E01), and no cross reactivity with c-peptide and known percentage to proinsulin (43% to mouse proinsulin I and 60% to mouse proinsulin II (Mercodia product information) and can be subtracted out for the Insulin Elisa (Cat# 10-1247-10).

#### Isolation of mouse islets

Pancreatic islets were isolated from mice at 24 weeks of age as previously described and following injection of collagenase into the common bile duct, excision and digestion of the pancreas, and retrieval of the islets using a Histopaque gradient (Stull et al., 2012).

#### Quantitative RT-PCR

Mouse islets, muscle, heart, liver, and hypothalamus were washed twice with PBS, and total RNA was extracted using Qiagen RNeasy Micro kit. Total RNA was reverse-transcribed at 37°C for 1 h using 15 μg random hexamers, 0.5 mM deoxynucleotide triphosphate, 5× first-strand buffer, 0.01 mM DTT, and 200 U of Moloney murine leukemia virus reverse transcriptase (Invitrogen) in a final reaction volume of 20 μL (Evans-Molina et al., 2007). The cDNA template was then diluted 1:5 with RNase-free water and subjected to real-time reverse-transcriptase quantitative PCR (RT-qPCR) using SensiFAST SYBR Lo-ROX kit and a QuantStudio 3 thermal cycler. Relative RNA levels were calculated against the constitutively expressed β-actin mRNA using the comparative C_T_ method, as described previously (Chakrabarti et al., 2002). Primer sequences are provided in Key Resources Table.

#### Mouse islet Ca^2+^ imaging

Direct measurement of ER Ca^2+^ was performed as previously described (Kono et al., 2018; Tong *et al*., 2016). Briefly, isolated mouse islets were incubated for 24 h with an adenovirus encoding the ER-targeted D4ER probe expressed under transcriptional control of the rat insulin promoter (Kono *et al*., 2018; Yamamoto *et al*., 2019). Transduced islets were incubated for an additional 24 h and transferred for imaging to a glass-bottom plate containing HBSS supplemented with 0.2% BSA, 1.0 mM Mg^2+^, and 2.0 mM Ca^2+^. For Förster Resonance Energy Transfer (FRET) experiments, confocal images were acquired with an LSM 800 confocal imaging system (Carl Zeiss). Imaging was performed using a 448 nm single excitation laser line and two hybrid fluorescence emission detectors set to 460-500 nm and 515-550 nm slit width. Time-series images (z-stacks) of 2-3 stage-registered fields were acquired over a period of 20 minutes. FRET ratios were calculated using the Zen Blue edition version 2.3 software.

For cytosolic Ca^2+^ imaging, islets were incubated in HBSS buffer supplemented with 2 mM Ca^2+^ and loaded with the ratiometric Ca^2+^ indicator fura-2-acetoxymethylester (Fura-2 AM). (Life Technologies). Baseline measurements were performed at 5.5 mM glucose and spontaneous intracellular Ca^2+^ transients were measured in response to 16.7 mM using a Zeiss Z1 microscope, as previously reported (Kono *et al*., 2018; Yamamoto *et al*., 2019).

To analyze calcium synchronicity, islets were transduced with a GCaMP6s expressing adenovirus and stimulated with high glucose (16.7 mM). Images were captured using a Zeiss LSM800 microscope. Islet movement was corrected by registration in ImageJ (Parslow et al., 2014). Regions of interest (ROI) were manually assigned to cells positive for GCaMP6s. The mean intensity of cell ROIs was measured in ImageJ for each cell over the period of duration of the experiment. The average of the mean intensities was calculated for representative islets and plotted versus time with the 95% confidence interval. The average GCaMP6s signal was calculated by taking the average fluorescent intensity of the cell over the course of the experiment (FI_avg_) for each islet. The Z-score was calculated for each time point and each β cell by subtracting the average fluorescent intensity of the β cell over the course of the experiment (FI_avg_) from the fluorescent intensity at each time point (FI_t_) and dividing by the standard deviation (StDevFI_avg_) of the β cell fluorescent intensity over the course of the experiment; Z-Score = (FI_t_ - FI_avg_) / StDevFI_avg_.

The Average Standard Deviation of Islet β Cell Z-Score was calculated for each islet by taking the average of the standard deviation of β cell Z-scores for each islet over the course of the experiment. Oscillations in calcium were illustrated by displaying the calculated Z-score of each β cell separated by islet with a red divider. Z-score values greater than 1.96 represent statistical elevations in GCaMP6s signal (alpha ≤ 0.05).

#### Immunoblotting

Immunoblotting was performed as previously described (Kono *et al*., 2012). Approximately 1.5 × 10^5^ S2KO and WT INS-1 cells or 100-125 human or mouse islets were incubated in a RPMI culture medium for 2 days (cells) or 1 day (islets). Mouse and human islets were treated with 0.5 mmol/L BSA-conjugated palmitate and 25 mmol/L glucose for 24 h to model glucolipotoxicity (GLT). S2KO and WT INS-1 cells were treated with either an adenovirus encoding human SERCA2b expressed for 24 h (Park et al., 2010), 4 µM brefeldin A for 24 h, or 10 µM cycloheximide (CHX) in the presence or absence of 10 µM MG-132 proteasome inhibitor for 9 h prior to protein extraction. Following the specified treatment, cells and islets were washed with PBS and lysed with buffer containing 50 mM Tris (pH 8.0), 150 mM NaCl, 0.05% deoxycholate, 0.1% IGEPAL CA-630 (Sigma-Aldrich, St. Louis, MO), 0.1% SDS, 0.2% sarcosyl, 10% glycerol, 1 mM dithiothreitol (DTT), 1 mM EDTA, 10 mM NaF, EDTA-free protease inhibitors, phosphatase inhibitors, 2 mM MgCl_2_, and 0.05% v/v benzonase nuclease. Mouse muscle, heart, and liver were also homogenized and lysed using the same buffer and assayed for SERCA2. Protein concentration was measured using the DC (detergent compatible) protein assay and a SpectraMax M5 multiwell plate reader. Equal quantity of proteins were suspended in 10% SDS and subjected to electrophoresis on a 4–20% Mini-Protean TGX gel in a Mini-Protean Tetra apparatus. Proteins were transferred to a PVDF membrane, and the membrane was blocked with Odyssey blocking buffer and incubated with primary antibodies in Signal Enhancer Hikari at 4°C overnight. Primary antibodies were diluted 1:1000, except β-actin, which was diluted 1:10,000. Bound primary antibodies were detected with anti-mouse donkey antibody (1:10,000 dilution), anti-goat donkey antibody (1:10,000 dilution), or anti-rabbit donkey antibody (1:10,000 dilution) (Kono *et al*., 2012). Immunoblots were scanned using LI-COR Biosciences Odyssey 1828 scanner and analyzed using Image Studio software (version 3.1.4).

#### Bulk RNA sequencing

Islets from three control and three βS2KO male mice were hand-picked and incubated overnight in regular RPMI supplemented with 10% FBS, 100 U/mL penicillin, and 100 μg/mL streptomycin. RNA was isolated the following day using the RNeasy Micro kit, and the quantity and quality of RNA were evaluated using the Bioanalyzer 2100. The cDNA library was prepared using 100 ng of total RNA and the KAPA mRNA Hyper Prep kit, following the manufacturer’s protocol. For each generated indexed library, the quantity and quality were assessed by Qubit and Bioanalyzer 2100, respectively. Multiple libraries were pooled in equal molarity. The pooled libraries were denatured, neutralized, and loaded to HiSeq 4000 sequencer at 300 pM final concentration for 100b paired-end sequencing. Approximately 30M reads per library were generated. The Phred quality score (Q score) was used to measure the quality of sequencing; more than 90% of the sequencing reads reached Q30 (99.9% base call accuracy). The quality of sequencing data was assessed using the FastQC tool.

#### mRNA sequencing data analysis

Sequencing files were analyzed using the Flow software. The raw sequencing files were aligned to the mouse genome (mm10) using STAR aligner. Uniquely mapped reads were annotated using RefSeq. mRNAs with a total of at least 10 read counts were retained for further analysis. mRNAs with linear scale fold-change ≥ 1.5 and p < 0.05 were identified using DESeq2 and were considered as differentially expressed mRNAs. Biological pathways were identified using QIAGEN Ingenuity Pathway Analysis, and gene ontology enrichment was performed using Metascape. Functional terms with p < 0.05 were considered significant. Figures were generated using the ggplot2 package and GraphPad.

#### Fluorogenic PC1/3 and PC2 enzyme activity assays

S2KO or WT INS-1 cells were transduced with an empty adenovirus or adenovirus expressing human SERCA2b (1 × 10^7^ PFU/well) (a kind gift from Dr. Umut Ozcan, Harvard Medical School) (Park *et al*., 2010). Forty-eight hours after transduction, cells were incubated for an additional 24 h in Opti-Mem I medium containing 11 mM glucose, 100 U/mL penicillin, 100 μg/mL streptomycin, 10 mM HEPES, 2 mM L-glutamine, 1 mM sodium pyruvate, 50 μmol/mL β-mercaptoethanol, and 1 mg/mL bovine aprotinin (Cayman Chemical). Enzyme activities of secreted PC1/3 and PC2 were analyzed in a reaction buffer containing 100 mM sodium acetate (pH 5.5 for PC1/3; pH 5.0 for PC2), 2mM CaCl_2_, 0.1% Triton X-100, 200 μM pERTKR-aminomethyl-coumarin, and proteinase inhibitor cocktail containing 1 μM pepstatin, 0.28 mM N-p-tosyl-L-phenylalanine chloromethyl ketone, 10 μM trans-epoxysuccinyl-L-leucylamido(4-guanidino) butane, and 0.14 mM Nα-tosyl-L-lysine chloromethyl ketone. To specifically measure the activity of PC1/3 and PC2, micromolar 7B2-CT peptide was used to inhibit PC2 activity in parallel reactions, while a similar concentration of the proSAAS-CT peptide was used to inhibit PC1/3 activity (Hoshino et al., 2011). Following incubation at 37°C for 3 h, the fluorescence was quantified using the SpectraMax ID5 plate reader operating at the excitation wavelength of 380 nm and the emission wavelength of 460 nm. Aminomethyl-coumarin was used as a standard (Apletalina et al., 1998; Blanco et al., 2015). Enzyme activity was normalized to the amount of the enzyme determined by immunoblotting; inhibitor-containing reactions were subtracted from non-inhibitor-containing reactions.

#### Immunofluorescence and morphometric analysis

Mice were euthanized and pancreata were rapidly removed and fixed overnight in Z-fix buffered zinc formalin. Fixed specimens were paraffin-embedded and longitudinal sections measuring 5 μm in thickness were obtained (Tong *et al*., 2016). β cell mass was estimated in each animal by multiplying the β cell fractional area by the weight of the pancreas, as previously described (Tong *et al*., 2016). After reducing non-specific binding with 30 min of Animal-Free Blocker incubation followed by 1 h incubation with Mouse on Mouse Blocking Reagent (Vector Laboratories), sections were incubated with primary antibodies (Proinsulin, Insulin, GM130, Giantin, Calnexin, LMAN1) at 4°C overnight in a humidity chamber and then with AlexaFluor-conjugated secondary antibodies at room temperature for 1 h. Stained sections were mounted with Fluorosave. Islet images were acquired using the LSM 800 confocal imaging system using 40X and 63X oil immersion objectives with an Airyscan detector. Images were analyzed using Zen Blue edition software. Signal intensities were quantified by ImageJ. Puncta count and colocalization analyses were performed using CellProfiler 4.1.3 (cellprofiler.org) (McQuin et al., 2018). Background was subtracted from each image by removing the lower-quartile intensity from each channel. Pancreas regions of interest (ROIs) were defined by proinsulin-positive area, and proinsulin, LMAN1, and Giantin puncta were identified by discarding objects outside a pixel diameter range after median filtering (Muralidharan et al., 2021). The puncta count was normalized to the proinsulin-positive area. For colocalization positivity, parent (Giantin or LMAN1) and child (proinsulin) objects were defined (Muralidharan *et al*., 2021), and the percentage of proinsulin puncta colocalized or co-compartmentalizing with Giantin or LMAN was determined using CellProfiler’s built in tutorials.

### QUANTIFICATION AND STATISTICAL ANALYSIS

The definitions of n, mean, and error bars, and the statistical methods utilized are listed in figure legends. Unless otherwise indicated, at least three independent experiments were carried out for each assay. Data are displayed as mean ± SEM, and a p-value of < 0.05 was considered to indicate a significant difference between groups. Data analysis was performed using the GraphPad Prism software. Pairwise comparisons were performed using unpaired Student’s *t-* test. Interactions between two groups with multiple treatments were determined using two-way ANOVA with a multiple comparisons post-hoc test. Normal distribution was assessed with a Kolmogorov-Smirnov test, and when the p-value was > 0.05, normal data distribution was assumed.

## Supporting information

Supplemental Tabale S2

## AUTHOR CONTRIBUTIONS

H.I. designed and conducted the experiments, analyzed the data, and wrote the manuscript. T.K. contributed to funding acquisition, conception and design of the study, data analysis and interpretation, collection and assembly of data, and manuscript writing/editing. C.L. (animal model), P.K. (Bulk RNA seq), M.C.A. (GCaMP6 analysis), S.A.W. (Imaging data analysis, manuscript writing/editing), T.J. (proinsulin localization), R.N.B. (ERGIC and Golgi co-localization analysis), and X.T. (in vitro model) participated in data collection and critical revision of the manuscript. P.A. and I.L. contributed to the research design, directed protein processing analysis experiments, provided critical reagents, and provided critical revision of the manuscript. C.E-M. directed funding acquisition, study conception and design, directed manuscript writing, and gave final approval of the manuscript. C.E-M. is the guarantor of this work, had full access to all of the study data, and takes responsibility for the integrity and accuracy of the data.

## ACKNOWLEDGEMENTS

The authors would like to thank Dr. Emily Anderson-Baucum for her helpful advice and edits. The authors also thank Kara Orr and Jacqueline Aquino (Indiana University) for their technical assistance.

---

**Supplemental Figure S1.**
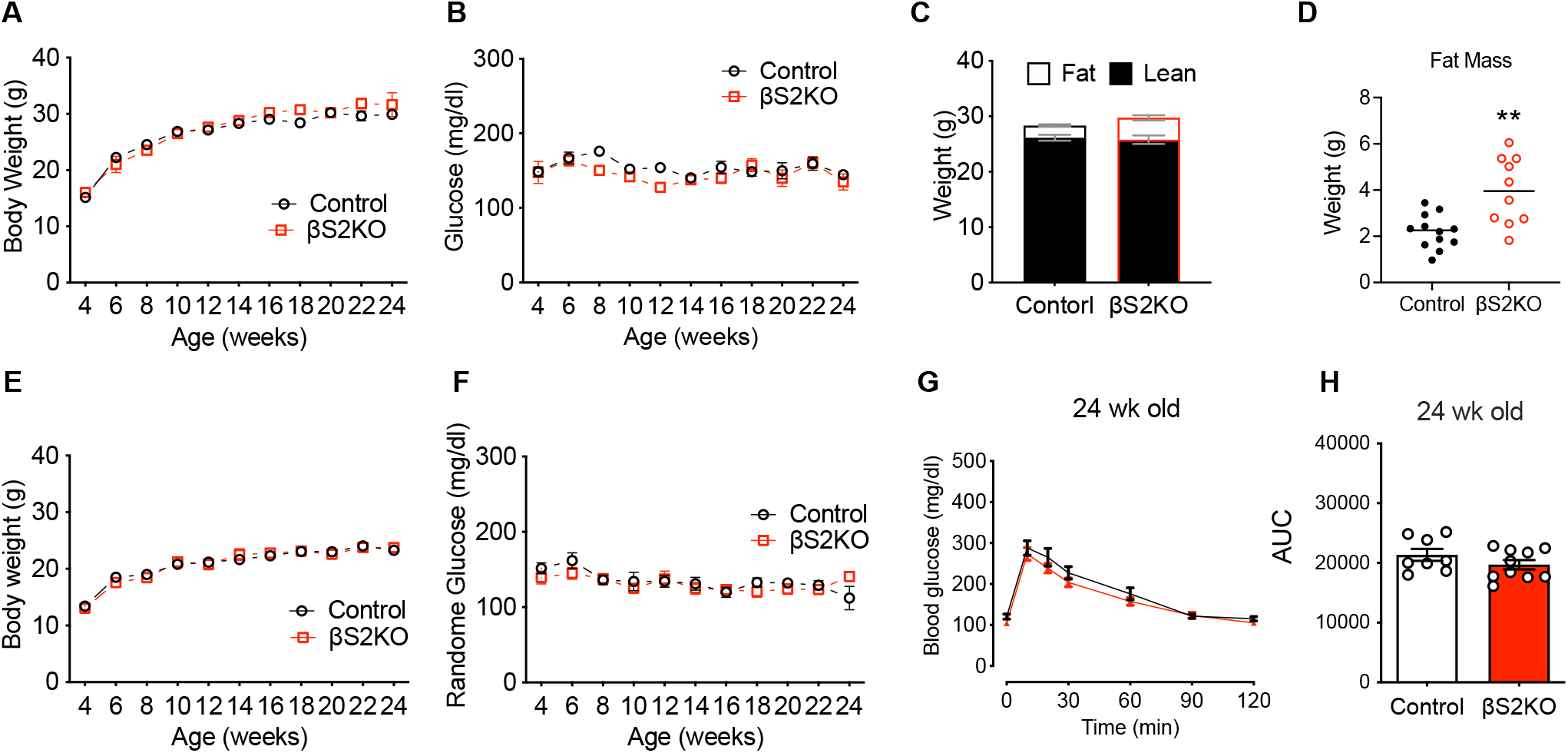
SERCA2 deficiency did not show any effects on weight change, lean mass, nor glucose tolerance in female mice. β cell-specific SERCA2 KO (βS2KO) and SERCA2_flox/flox_ mice (Control) were fed a normal chow diet for 25 weeks. Longitudinal changes in whole body weight and random blood glucose in male (A and B) and female mice (E and F) was monitored until 24-week-old, n= at least 6. (C-D) Lean mass and fat mass were measured during the experimental period using the EchoMRI 500 Body Composition Analyzer, n=10-12. (G-H) GTT was performed (2 g/kg glucose dosed to lean mass) in female mice at 24 weeks of age. Area under curve (AUC) analysis is shown graphically (H), n=8-10. Results are displayed as mean ± SEM. *p < 0.05; **p < 0.01, vs control.

**Supplemental Figure S2.**
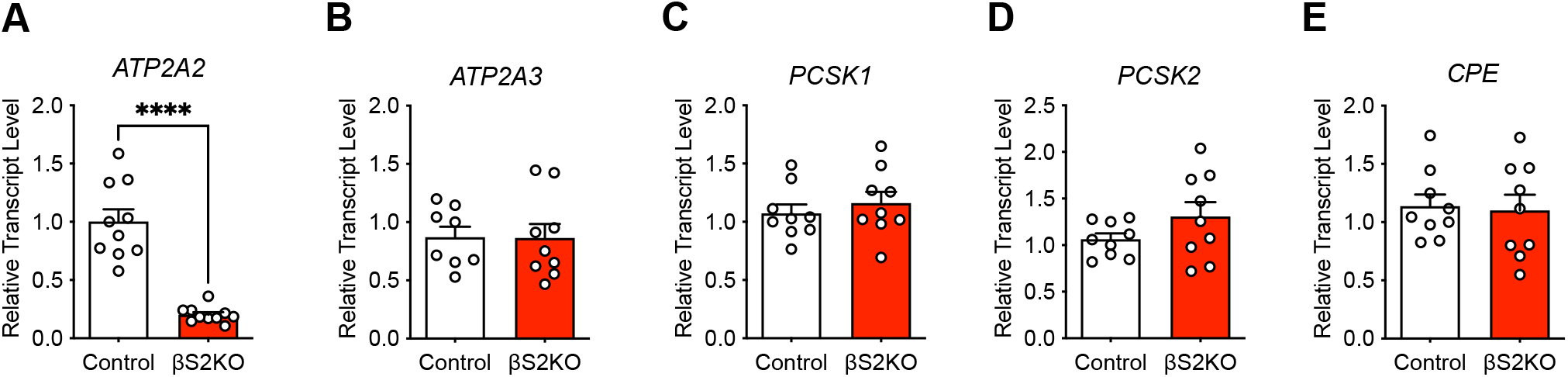
Transcript levels of *ATP2A2, ATP2A3, PCSK1, PCSK2, CPE* in islets of control and βS2KO mice (male, 24 weeks of age) were determined by RT-qPCR and normalized to actin levels. Replicates are indicated by open circles and squares. *Indicates statistically significant difference from control (****p<0.0001) by a Student’s *t*-test.

**Supplemental Figure S3.**
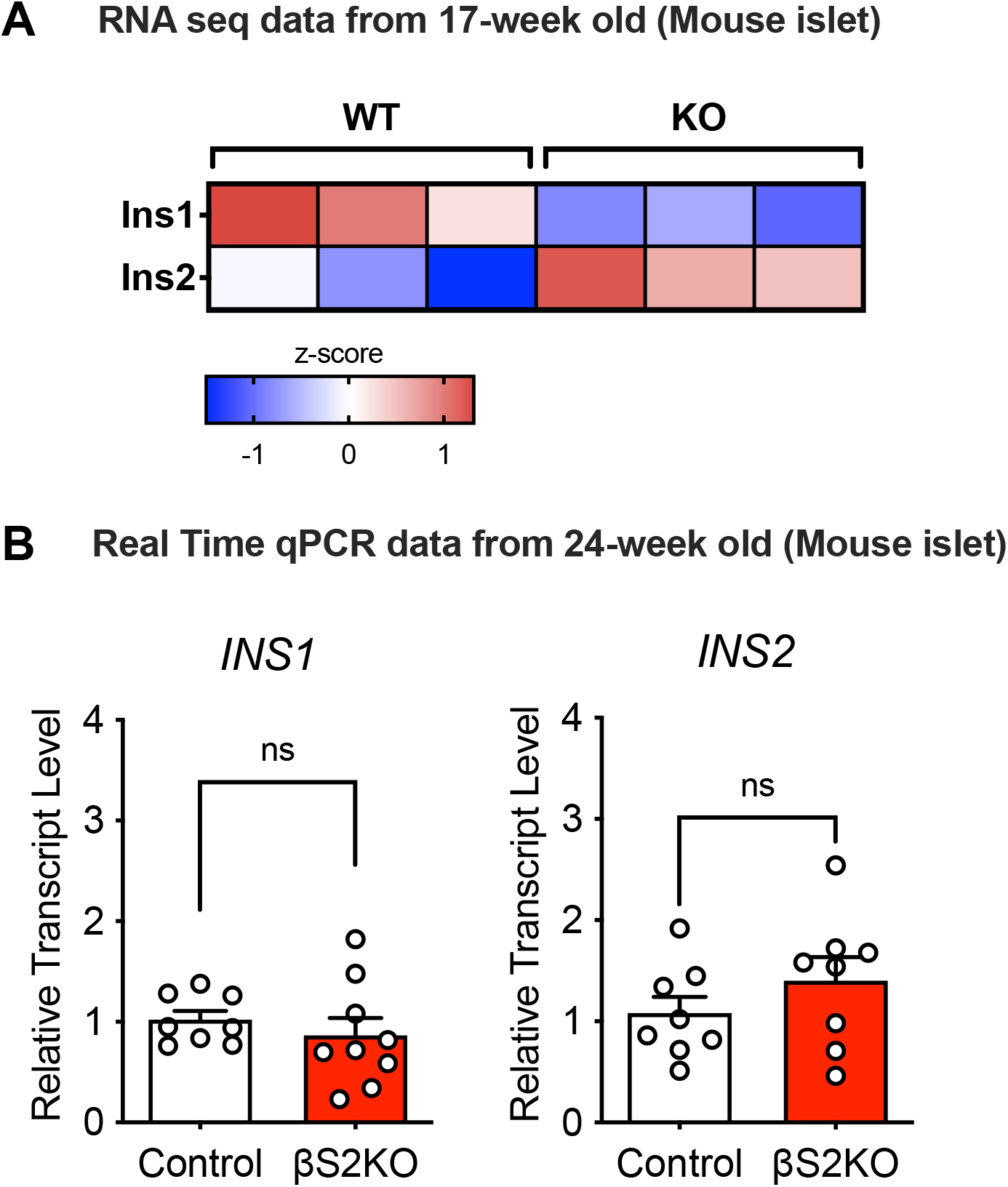
**(A)** RNA isolated from control and βS2KO islets was subjected to bulk RNA sequencing analysis at 17-weeks of age. Heatmap of the fold changes for insulin genes. **(B)** Transcript levels of *INS1* and *INS2* in islets of control and βS2KO mice (male, 24 weeks of age) were determined by RT-qPCR and normalized to actin levels. Replicates are indicated by open circles and squares. ns = not significantly different between groups.

