## Supplemental Table S1 for "SERCA2 regulates proinsulin processing and processing enzyme maturation in the pancreatic β cell"

**KEY RESOURCES TABLE**

| REAGENT or RESOURCE | SOURCE | IDENTIFIER |
| --- | --- | --- |
| Antibodies | | |
| Rabbit polyclonal anti-PC1/3 | Cell Signaling Technology | Cat# 11914; RRID:AB_2631284) |
| Rabbit polyclonal PC1/3 C-terminal | Gift from Dr. Iris Lindberg, University of Maryland  Vindrola et al., 1992 | http://thelindberglab.com/antibodies/ |
| Rabbit monoclonal anti-PC2 | Cell Signaling Technology | Cat# 14013; RRID:AB_2631285 |
| Rabbit polyclonal pro-PC2 | Gift from Dr. Iris Lindberg, University of Maryland  Muller et al., 2000 | http://thelindberglab.com/antibodies/ |
| Rabbit polyclonal anti-CPE | Abcam | Cat# ab11044; RRID:AB_297698 |
| Goat polyclonal anti-SERCA2 | Santa Cruz Biotechnology | Cat# sc-8095; RRID:AB_2290108 |
| Mouse monoclonal anti-Actin | Millipore | Cat# MAB1501; RRID:AB_2223041 |
| Guinea pig polyclonal anti-Insulin | Agilent Dako | Cat# A0564; RRID:AB_10013624 |
| Mouse monoclonal anti-proInsulin | Developmental Studies Hybridoma Bank | Cat# GS-9A8-C |
| Rabbit monoclonal anti-Insulin (C27C9) | Cell Signaling Technology | Cat# 3014, RRID:AB_2126503 |
| Rabbit monoclonal anti-LMAN1 (ERGIC53) | Abcam | Cat# ab125006; RRID:AB_10973984 |
| Goat polyclonal Anti-Giantin | Santa Cruz Biotechnology | Cat# sc-46993; RRID:AB_2279271 |
| IIRDye® 800CW Donkey anti-Mouse IgG Secondary Antibody | LI-COR Bioscience | Cat# 926-32212, RRID:AB_621847 |
| IRDye® 800CW Donkey anti-Goat IgG Secondary Antibody | LI-COR Bioscience | Cat# 926-32214, RRID:AB_621846 |
| IRDye® 680RD Donkey anti-Rabbit IgG Secondary Antibody | LI-COR Bioscience | Cat# 926-68073, RRID:AB_10954442 |
| Donkey anti-guinea pig IgG Secondary Antibody (Alexa Fluor-647) | Jackson ImmunoResearch Laboratories, Inc | **Cat# 706-605-148, RRID:** AB_2340476 |
| Donkey anti-goat IgG Secondary Antibody (Alexa Fluor-568) | Thermo Fisher Scientific | Cat# **A11057, RRID:** AB_ 2534104 |
| Donkey anti-rabbit IgG Secondary Antibody (Alexa Fluor-647) | Thermo Fisher Scientific | Cat# A31573**, RRID:** AB_2536183 |
| Donkey anti-mouse IgG Secondary Antibody (Alexa Fluor-488) | Thermo Fisher Scientific | Cat# A-21202**, RRID:** AB_141607 |
| Mouse-on-Mouse IgG Blocking Solution | Vector Laboratories | Cat# MKB-2213-1 |
| Vector® NovaRED™ Substrate Kit, Peroxidase | Vector Laboratories | Cat# SK-4800 |
| Bacterial and virus strains | | |
| adenovirus expressing human SERCA2b | Gift from Dr. Umut Ozcan, Harvard Medical School | N/A |
| adenovirus expressing RIP-D4ER Cameleon probe | Gift from Dr. Richard Benninger, University of Colorado  Greotti et al., 2016 | N/A |
| Adenovirus expressing RIP-GCaMP6s | VectorBuilder | N/A |
| Biological samples | | |
| Human donor islets | See Supplemental Table S2 | N/A |
| Chemicals, peptides, and recombinant proteins | | |
| Insulin (regular, short-acting) | Novo Nordisk | NC0769896 |
| D-Glucose | Sigma Aldrich | Cat# G7528 |
| Brefeldin A (BFA) | Sigma Aldrich | Cat# B6542 |
| Palmitate | Sigma Aldrich | Cat# P0500 |
| Bovine aprotinin | Cayman Chemical | Cat# 14716 |
| Cyclohexamide | Sigma Aldrich | Cat# C4859 |
| MG132 | Sigma Aldrich | Cat# M7449 |
| RPMI1640 | Gibco | Cat# 11875-093 |
| pERTKR-aminomethylcumarin | Peptide International | Cat# MPR-3159-v |
| Aminomethylcumarin | Sigma Aldrich | Cat# A9891 |
| N-p-tosyl-L-phenylalanine chloromethyl ketone | Sigma Aldrich | Cat# T4376 |
| trans-epoxysuccinyl-L-leucylamido(4-guanidino) butane | Sigma Aldrich | Cat# 66701 |
| Nα-tosyl-L-lysine chloromethyl ketone | Sigma Aldrich | Cat# T7254 |
| 7B2-CT peptide | Gift from Dr. Iris Lindberg, University of Maryland | N/A |
| ProSAAS-CT peptide | Gift from Dr. Iris Lindberg, University of Maryland | N/A |
| Critical commercial assays | | |
| RNeasy Mini Kit | Qiagen | Cat# 74136 |
| RNeasy Micro Kit | Qiagen | Cat# 74034 |
| DNeasy Blood and Tissue kit | Qiagen | Cat# 69504 |
| SensiFASTÔSYBR Lo-ROX kit | Bioline | Cat# BIO-94020 |
| BCA protein determination | BioRad | Cat# 500-0112 |
| Mouse Insulin ELISA | Mercodia | Cat# 10-1247-10 |
| Fura-2 acetoxymethylester | Invitrogen | Cat# F1221 |
| Mouse ProInsulin ELISA | Alpco Diagnostics | Cat# 80-PINMS-E01 |
| KAPA mRNA Hyper Prep Kit | Roche | Cat# KK8540 |
| Experimental models: Cell lines | | |
| Rat (male) SERCA2 null INS-1 832/13 | This paper | N/A |
| Rat (male) INS-1 832/13 | H E Hohmeier et al., 2000 | N/A |
| Experimental models: Organisms/strains | | |
| Mouse: β cell specific SERCA2-null (βS2KO) in C57BL6/J background | This paper | N/A |
| Mouse: C57BL6/J | The Jackson Laboratory | JAX # 000664 |
| Oligonucleotides | | |
| Mouse PC1/3-F | This paper | AGTTGGAGGCATAAGAATGCTG |
| Mouse PC1/3-R | This paper | GCCTTCTGGGCTAGTCTGC |
| Mouse PC2-F | This paper | AGAGAGACCCCAGGATAAAGATG |
| Mouse PC2-R | This paper | CTTGCCCAGTGTTGAACAGGT |
| Mouse CPE-F | This paper | GCTCAGGTAATTGAAGTCTT |
| Mouse CPE-R | This paper | TACTGCTCACGAATACAGTT |
| Mouse SERCA2b-F | This paper | GATCCTCTACGTGGAACCTTTG |
| Mouse SERCA2b-R | This paper | CCACAGGGAGCAGGAAGAT |
| Mouse SERCA3-F | This paper | AGGGGAAGCTAAGAAGCCAG |
| Mouse SERCA3-R | This paper | CCCTCAGACTCCTCCTACCC |
| Mouse beta-actin-F | This paper | AGGTCATCACTATTGGCAACGA |
| Mouse beta-actin-R | This paper | CACTTCATGATGGATTGAATGTAGTT |
| Software and algorithms | | |
| Zen Blue edition ver2.3 | Carl Zeiss | [https://www.zeiss.com/microscopy/int/products/microscope-software.html](about:blank); RRID:SCR_013672 |
| Axio-Vision Software | Carl Zeiss | [https://www.zeiss.com/microscopy/int/products/microscope-software.html](about:blank); RRID:SCR_002677 |
| ImageJ | Fiji; Schneider et al., 2012 | Open source: [https://imagej.net/Fiji](about:blank); RRID:SCR_002285 |
| Image Studio | LI-COR | [https://www.licor.com/bio/image-studio-lite/download](about:blank); RRID:SCR_013715 |
| FastQC | Babraham Bioinformatics | [https://www.bioinformatics.babraham.ac.uk/projects/download.html](about:blank)  RRID:SCR_011106 |
| Prism 7.0 | GraphPad Software | [https://www.graphpad.com/](about:blank); RRID:SCR_002798 |
| Flow software version 10.0.20.1231 | Partek | [https://www.partek.com/partek-flow/](about:blank); RRID:SCR_011860 |
| STAR aligner ver. 2.7.3a | Dobin A et al., 2013 | [https://github.com/alexdobin/STAR/releases](about:blank); RRID:SCR_004463 |
| RefSeq release 93 | NCBI | [https://www.ncbi.nlm.nih.gov/refseq/](about:blank); RRID:SCR_003496 |
| DESeq2 | Bioconductor | Open source: [https://bioconductor.org/packages/release/bioc/html/DESeq2.html](about:blank); RRID:SCR_015687 |
| Ingenuity Pathway Analysis | Qiagen | [https://digitalinsights.qiagen.com/product-login/](about:blank); RRID:SCR_008653 |
| Metascape | Metascape | [https://metascape.org](about:blank); RRID:SCR_016620 |
| ggplot2 | MIT | [https://ggplot2.tidyverse.org](about:blank); RRID:SCR_014601 |
| Other | | |
| Perifusion System | Biorep Technologies, Inc | N/A |
| LSM 800 confocal imaging system | Carl Zeiss | RRID:SCR_015963 |
| LSM-700 confocal microscope | Carl Zeiss | RRID:SCR_017377 |
| Bioanalyzer 2100 | Agilent | Cat# G2939BA; RRID:SCR_019715 |
| HiSeq 4000 sequencer | Illumina | RRID:SCR_016386 |
| Odyssey CLx scanner | LI-COR | RRID:SCR_014579 |
| EchoMRI-500 | EchoMRI | RRID:SCR_017104 |
| Contour | Bayer | N/A |
